# SiCloneFit: Bayesian inference of population structure, genotype,and phylogeny of tumor clones from single-cell genome sequencing data

**DOI:** 10.1101/394262

**Authors:** Hamim Zafar, Nicholas Navin, Ken Chen, Luay Nakhleh

## Abstract

Accumulation and selection of somatic mutations in a Darwinian framework result in intra-tumor heterogeneity (ITH) that poses significant challenges to the diagnosis and clinical therapy of cancer. Identification of the tumor cell populations (clones) and reconstruction of their evolutionary relationship can elucidate this heterogeneity. Recently developed single-cell DNA sequencing (SCS) technologies promise to resolve ITH to a single-cell level. However, technical errors in SCS datasets, including false-positives (FP), false-negatives (FN) due to allelic dropout and cell doublets, significantly complicate these tasks. Here, we propose a non-parametric Bayesian method that reconstructs the clonal populations as clusters of single cells, genotypes of each clone and the evolutionary relationships between the clones. It employs a tree-structured Chinese restaurant process as the prior on the number and composition of clonal populations. The evolution of the clonal populations is modeled by a clonal phylogeny and a finite-site model of evolution to account for potential mutation recurrence and losses. We probabilistically account for FP and FN errors, and cell doublets are modeled by employing a Beta-binomial distribution. We develop a Gibbs sampling algorithm comprising of partial reversible-jump and partial Metropolis-Hastings updates to explore the joint posterior space of all parameters. The performance of our method on synthetic and experimental datasets suggests that joint reconstruction of tumor clones and clonal phylogeny under a finite-site model of evolution leads to more accurate inferences. Our method is the first to enable this joint reconstruction in a fully Bayesian framework, thus providing measures of support of the inferences it makes.

## Introduction

Acquisition of somatic mutations that confer selective growth advantage to the carrier cells drives initiation and progression of cancer (Vogelstein et al., 2013). From an evolutionary viewpoint, tumor progression is a somatic evolutionary process that gives rise to a composite mixture of genetically distinct subpopulations (clones) of cells through rounds of accumulation of somatic alterations, proliferation and Darwinian selection in the tumor microenvironment (Nowell, 1976; Merlo et al., 2006; Pepper et al., 2009; Yates and Campbell, 2012). The genomic heterogeneity within a tumor, also known as intra-tumor heterogeneity (ITH) not only propels disease progression and metastasis (Turke et al., 2010; Wu et al., 2012), but can also lead to therapeutic relapse and drug resistance (Gillies, Verduzco, and Gatenby, 2012; Burrell et al., 2013). High-throughput next-generation sequencing (NGS) technologies have provided large-scale quantitative genomic datasets (Nik-Zainal et al., 2012; Kandoth et al., 2013) for investigating ITH. Most studies typically perform deep sequencing of bulk DNA retrieved from a single sample of the cancer tissue (Shah et al., 2012; Landau et al., 2015). Such datasets provide variant allele frequencies (VAFs) of somatic mutations, an aggregate signal averaged over the existing distinct tumor subclones as well as contaminating normal cells (Navin, 2014), and VAFs are modeled as mixtures of subclones for their computational inference (Roth et al., 2014; Deshwar et al., 2015; El-Kebir et al., 2016; Jiang et al., 2016). However, noisy aggregate signal of VAFs has limited resolution and thus restricts a comprehensive exploration of ITH (Navin, 2014; Baslan and Hicks, 2017). Sequencing multiple samples from different geographical regions of a tumor can improve upon single-sample bulk sequencing (Gerlinger et al., 2012; Gerlinger et al., 2014; Yates et al., 2015) but cannot resolve spatially intermixed subpopulations (Navin, 2015).

Ultimately, a single cell is the fundamental substrate of tumor evolution and single-cell DNA sequencing (SCS) has emerged as a powerful technique for resolving tumor evolution and ITH to a single-cell level (Hou et al., 2012; Wang et al., 2014; Gawad, Koh, and Quake, 2014; Leung et al., 2017). Such technologies provide sequencing data pertaining to single cells, thus allowing for direct measurement of genotypes and prevalences of tumor subclones without requiring deconvolution of aggregate signals (Zafar et al., 2018). At the same time, they offer the possibility of reconstructing the clonal lineage tree. However, these tasks are challenged by high level of experimental noise introduced in SCS data (Zafar et al., 2018) during the sample preparation and whole genome amplification (WGA) steps. WGA errors include: false-positive (FP) and false-negative (FN) errors due to allelic dropout (ADO) (Navin, 2014). FP errors are caused by deamination of cytosine bases and infidelity of polymerase enzymes. ADO affects the heterozygous loci as one of the alleles is preferentially amplified. Unintended isolation and processing of two cells together can result in cell doublets (characterized by merged genotype) (Zafar et al., 2018). Another problem with SCS data is missing entries due to coverage non-uniformity (Zafar et al., 2018).

Single-cell somatic point mutation profiles have been used to infer clonal subpopulations. Early studies (Wang et al., 2014; Li et al., 2012) used multidimensional scaling and hierarchical clustering for reconstructing the tumor subclones but such approaches fail to account for errors. Gawad *et al.*(Gawad, Koh, and Quake, 2014) used a Bernoulli mixture model (BMM) to infer clusters of cells and predict cluster genotypes and performed model selection via a Bayesian information criterion (BIC) score. This approach was extended in the SCG method (Roth et al., 2016) to accommodate errors due to ADO and doublets. However, such approaches neither utilize the evolutionary relationship between the clonal clusters nor infer any phylogeny that can convey the evolutionary history of the tumor cells. Another direction with SCS data has been the reconstruction of cell lineages to study tumor evolution. SCITE (Jahn, Kuipers, and Beerenwinkel, 2016) and OncoNEM (Ross and Markowetz, 2016) probabilistically model WGA-specific errors for inferring tumor lineages from SCS data. However, both SCITE and OncoNEM operate under the infinite sites assumption (ISA), which posits that no genomic site mutates more than once and mutations are never lost. This assumption could get violated in tumor evolution due to events including: convergent evolution, chromosomal deletions and loss of heterozygosity (LOH) (Davis and Navin, 2016; Kuipers et al., 2017). SiFit (Zafar et al., 2017) employs a finite-site model of evolution to allow for mutation recurrence and losses and employs a maximum-likelihood based approach for reconstructing tumor phylogeny. However, these phylogeny approaches (other than OncoNEM) do not provide straight-forward reconstruction of the tumor subclones. At the same time, none of these phylogeny-based methods account for cell doublets as the merged genotypes can not be represented by a cell lineage tree model.

Here, we propose SiCloneFit, a unified statistical framework and computational method that simultaneously addresses the problems of subclonal reconstruction and phylogeny inference from single-cell sequencing data. Our unified model simultaneously (i) estimates the number of tumor clones, (ii) identifies the tumor clones as clusters of single cells, (iii) predicts the mutations associated with each tumor clone (clonal genotype), and (iv) under a finite-site model of evolution places the tumor clones at the leaves of a phylogenetic tree (clonal tree) that models their genealogical relationships. In doing so, the SiCloneFit model integrates non-parametric Bayesian mixture modeling based on a Chinese restaurant process with the finite-sites-based phylogenetic approach introduced in SiFit (Zafar et al., 2017). Using single-cell somatic point mutation profiles as input, SiCloneFit introduces a non-parametric Bayesian mixture model based on a phylogeny-based Chinese restaurant process (clusters reside at the leaves of a phylogeny) to identify clusters (clone) of cells that share mutations and resolves the clonal genotypes (mutations associated with a clonal cluster). The evolution of the clonal genotypes is modeled using a clonal phylogeny and a finite-site model of evolution that accounts for the effects of deletion, LOH and point mutations at the genomic sites. SiCloneFit adopts the probabilistic error model of SiFit to account for FP and FN errors in SCS. The doublet-aware model of SiCloneFit employs a Beta-binomial distribution to accommodate for the presence of cell doublets and augments the non-parametric Bayesian mixture model with another finite mixture model to allow for the placement of a potential doublet in two clonal clusters. We develop a Gibbs sampling algorithm comprised of partial reversible-jump and partial Metropolis-Hastings updates to explore the joint posterior space of all parameters. Through simulations, we show the superiority of our method compared to existing subclonal reconstruction methods under a wide variety of parameter settings. We finally applied SiCloneFit on experimental SCS datasets and simultaneously reconstructed clonal populations, clonal genotypes and clonal phylogeny. Joint inference of clonal populations and their genealogical relationships by SiCloneFit led to an improvement in resolving ITH for the experimental SCS datasets compared to the existing methods that treat the problems separately. To the best of our knowledge, SiCloneFit is the first Bayesian framework that jointly reconstructs clonal populations and their evolutionary history from SCS datasets under a finite-site model of evolution while accounting for cell doublets along with other WGA artifacts. The method is publicly available at https://bitbucket.org/hamimzafar/siclonefit.

## Results

### Overview of SiCloneFit Model

We start with a brief description of the formulation of the joint inference problem and the SiCloneFit model. Overview of the SiCloneFit model is given in Fig. 1a.

**Figure 1:**
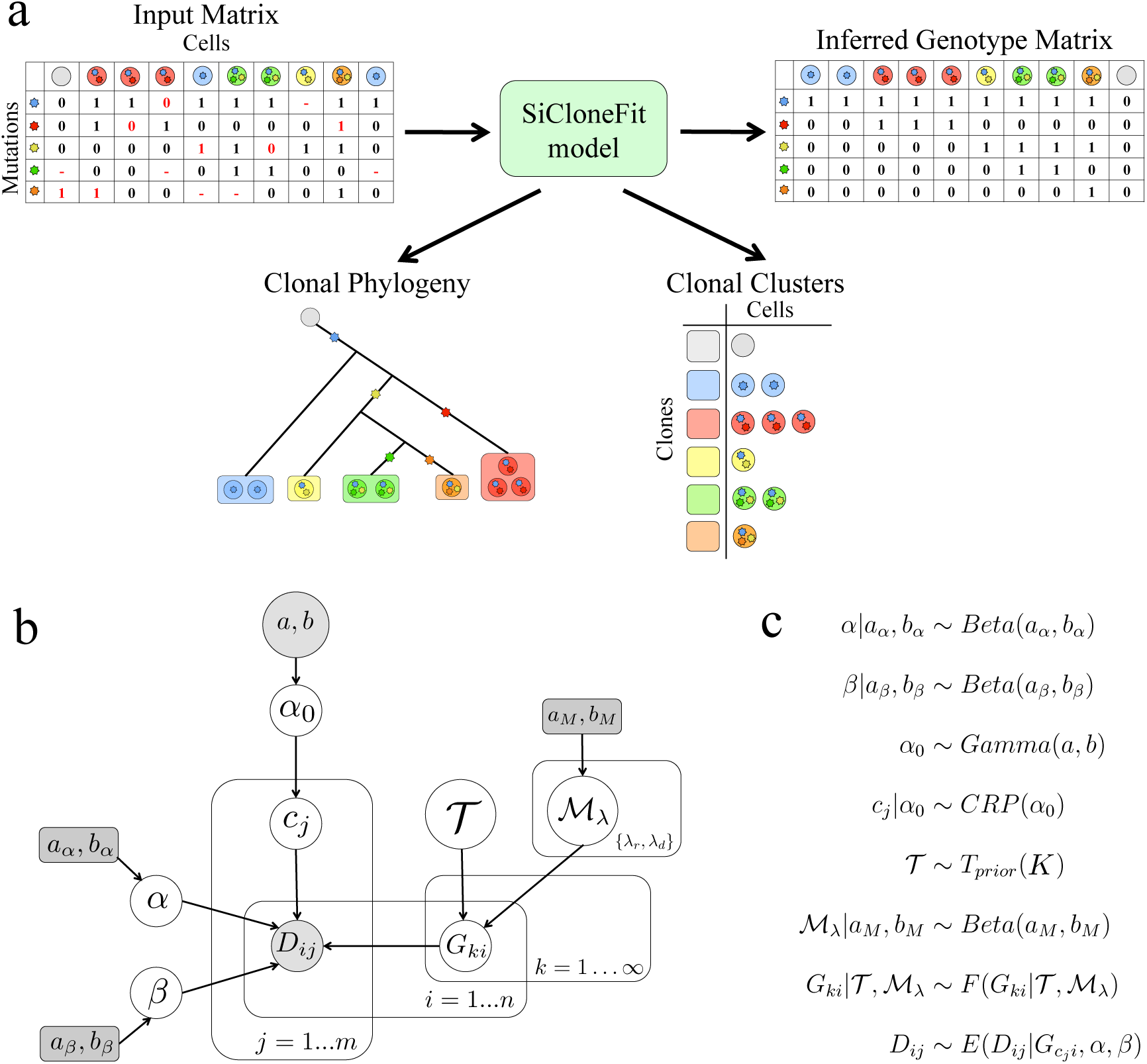
Overview of SiCloneFit Model. (a) From an observed noisy genotype matrix of single cells, SiCloneFit infers the clonal clusters, clonal phylogeny and clonal genotypes of single cells. (b) A probabilistic graphical model representing the singlet model of SiCloneFit. Shaded nodes represent the observed values or fixed parameters; unshaded nodes are the latent variables that are of interest; a posterior distribution over the values of the unshaded nodes is approximated using samples from the proposed Gibbs sampler. The variables and indices are described in Supplemental Methods. (c) Distributional assumptions for the different variables in the SiCloneFit singlet model.

A tumor population (clone) refers to a set of cells that share a common genotype as they descend from a common ancestor (Merlo et al., 2006). In the context of single-cell sequencing, a clonal population refers to a maximal set of cells with identical genotype (with respect to the set of mutations under analysis) (Roth et al., 2016). We model the lineage of the clonal populations using a clonal phylogeny, a rooted directed binary tree, the root of which represents normal (unmutated) genotype and somatic mutations are accumulated along the branches of the phylogeny. The sampling of single cells from the tumor at any point in time is analogous to horizontally slicing the clonal phylogeny to obtain samples from the leaves. The leaves of the clonal phylogeny represents the clonal populations and the sampled cells are individuals sampled from each leaf. The DNA from each sampled cell goes through the process of single-cell DNA sequencing and mutation calling, which provides the observed genotype matrix **D** = *D*_*n×m*_ for *m* single cells and *n* somatic mutation sites.

In SiCloneFit, we model this generative process using the probabilistic graphical model shown in Fig. 1b (also Supplemental Fig. S1). Here, we briefly describe the singlet model (all sampled cells are assumed to be singlets) of SiCloneFit. The probabilistic graphical model for the doublet-aware model is shown in Supplemental Fig. S2. The model variables, hyper-parameters and associated indices are introduced in Supplemental tables S1-S3. For a detailed description of the singlet and doublet-aware model of SiCloneFit, see Supplemental Methods.

We consider somatic single nucleotide variant (SNV) sites, where the input data is represented by a matrix that records the observed genotype for each cell for each mutations sites. The input matrix can be binary, when the presence or absence of a mutation is noted. For a ternary matrix, the three possible genotype states correspond to homozygous reference, heterozygous and homozygous non-reference genotypes. We assume that there is a set of *K* clonal populations from which a total of *m* single cells are sampled and the clonal populations can be placed at the leaves of a clonal phylogeny, *𝒯*. Each clonal population contains a set of cells that have identical genotype and share a common ancestor. It is important to note that *K* is unknown. To infer the number of clones and assign the cells to clones, we introduce a tree-structured infinite mixture model. In our model, we extend the tree-structured Chinese restaurant process (CRP) prior from (Meeds et al., 2008) to define a nonparametric Bayesian prior over binary trees, leaves of which represent the mixture components (clonal clusters). The clonal phylogeny represents the genealogical relationship between the clonal populations. The genotype vector associated with a clone is called clonal genotype and it records the genotype values for all mutation sites for the corresponding clone. To model the evolution of the clonal genotypes along the branches of *𝒯*, we employ a finite-site model of evolution, *ℳ*_*λ*_, that accounts for the effects of point mutations, deletion and LOH on the clonal genotypes. The model of evolution assigns transition probabilities to different genotype transitions along the branches of the clonal phylogeny. The true genotype of each cell is identical to the clonal genotype of the clonal cluster where it is assigned. However, observed genotypes of single cells can differ from their true genotype due to amplification errors introduced during the SCS work flow. The effect of amplification errors is modeled using an error model distribution parameterized by FP error rate *α* and FN error rate *β*. The generative process is described in detail in Methods and the distributional assumptions of the model are shown in Fig. 1c.

SiCloneFit attempts to jointly reconstruct the tumor clones as clusters of single cells, clonal genotypes and the clonal phylogeny. In doing so, it employs a likelihood function and a compound prior to define the posterior distribution over these latent variables. SiCloneFit employs a Markov chain Monte Carlo (MCMC) sampling procedure based on the Gibbs sampling algorithm comprised of partial reversible-jump and partial Metropolis-Hastings updates to estimate the latent variables. The posterior distribution and the inference algorithm are described in Methods and Supplemental Methods.

### Benchmarking on Simulated Datasets

We performed comprehensive simulations to evaluate the performance of SiCloneFit in (i) clustering the cells into different clones, (ii) inferring the genotypes of the cells via clonal genotyping and (iii) reconstructing the clonal lineage. To generate benchmarking datasets, we first sampled observed clonal prevalences for a fixed number of clones from a Dirichlet distribution and the cells were assigned to different clones using a multinomial distribution. Then we constructed linear and branching topologies for clonal phylogeny using the Beta-splitting model (Sainudiin and Veber, 2016). The clonal genotypes at the leaves of the phylogeny were simulated in a similar fashion as described in (Zafar et al., 2017). Different SCS artifacts were then introduced on the cellular genotypes to produce the noisy observed genotypes which were used as the input data for inference. The simulation process is described in detail in Supplemental Results.

To compare the results of SiCloneFit against the ground truth, we summarized the posterior samples from the Gibbs sampler of SiCloneFit. The clustering samples were summarized by the maximum posterior expected adjusted rand (MPEAR) method (Fritsch and Ickstadt, 2009). To summarize the clonal phylogeny samples, we constructed a maximum clade credibility topology (MCCT) from the posterior samples using *DendroPy* (Sukumaran and Holder, 2010). From the posterior samples, we computed the posterior probability of the genotype of each cell at each site and the genotype with the highest posterior probability was assigned as the inferred genotype. When using the doublet-aware model of SiCloneFit, the doublets were inferred based on the posterior probability and were filtered out for subsequent analysis. The summarization methods are described in detail in Supplemental Results.

We compared SiCloneFit’s performance against SCG (Roth et al., 2016) and OncoNEM (Ross and Markowetz, 2016). SCG was used to infer clonal genotypes and clonal structures from single cell somatic SNV profiles. The clonal phylogeny was obtained by running maximum parsimony algorithm (Schliep, 2011) on the clonal genotypes as suggested in (Roth et al., 2016). OncoNEM was used to infer a clonal tree from single cell somatic SNV profiles. Clonal genotypes were obtained by inferring the occurrence of the mutation on the branches of the clonal tree. The clustering accuracy of each method was measured using adjusted rand index and B-cubed F-score (Amigó et al., 2009) for datasets without and with doublets respectively. The genotyping performance was measured using Hamming distance (number of entries differing) between the true and inferred genotypes. For phylogeny inference, we used pairwise cell shortest-path distance (Ross and Markowetz, 2016) as the tree reconstruction error. The performance metrics are described in detail in Supplemental Results.

To evaluate SiCloneFit’s singlet model, we first ran simulations excluding doublets. For a fixed number of clones, we simulated datasets with varying numbers of cells and varying numbers of sites. For smaller sized (*m* = 100) datasets, we compared against SCG and OncoNEM, whereas, for larger sized datasets (*m* = 500), only SCG was compared as OncoNEM failed to run. Clustering accuracy (Fig. 2a, Supplemental Fig. S3) and phylogeny inference accuracy (Fig. 2c, Supplemental Fig. S5) of each method improved as the number of sites increased. Total genotyping error (Fig. 2b, Supplemental Fig. S4) increased with an increase in the number of sites. For each experimental setting, SiCloneFit performed the best in terms of all performance metrics. For larger sized datasets, it achieved perfect clustering for almost all datasets. In the presence of higher numbers of clonal populations, sampling the same number of cells leads to a more difficult inference problem. Even for such situations, SiCloneFit performed the best based on all three metrics and it was more robust against the increase in number of clones as evidenced by lower rate of reduction in clustering accuracy compared to SCG and OncoNEM (Supplemental Fig. S6). For larger datasets, we also tested the effect of missing data on inference accuracy. Even in the presence of high amount of missing data, SiCloneFit performed well in clustering the cells into clones and inferring the clonal phylogeny. It consistently performed better than SCG (Supplemental Fig. S7-S9) in terms of all metrics. Only in one setting (*n* = 100, 30% missing data) did SCG achieve lower genotyping error than SiCloneFit. SiCloneFit’s performance was also more robust against increasing error rate. With an increase in the FN rate, performance of each method degraded, but SiCloneFit had the lowest amount of reduction in performance and it also outperformed all the other methods for all values of false negative rate (Supplemental Fig. S10). Same trend was observed when FP rate was increased (Supplemental Fig. S11). In this setting, for some datasets, SCG’s genotyping failed to converge and resulted in a large number of false predictions.

**Figure 2:**
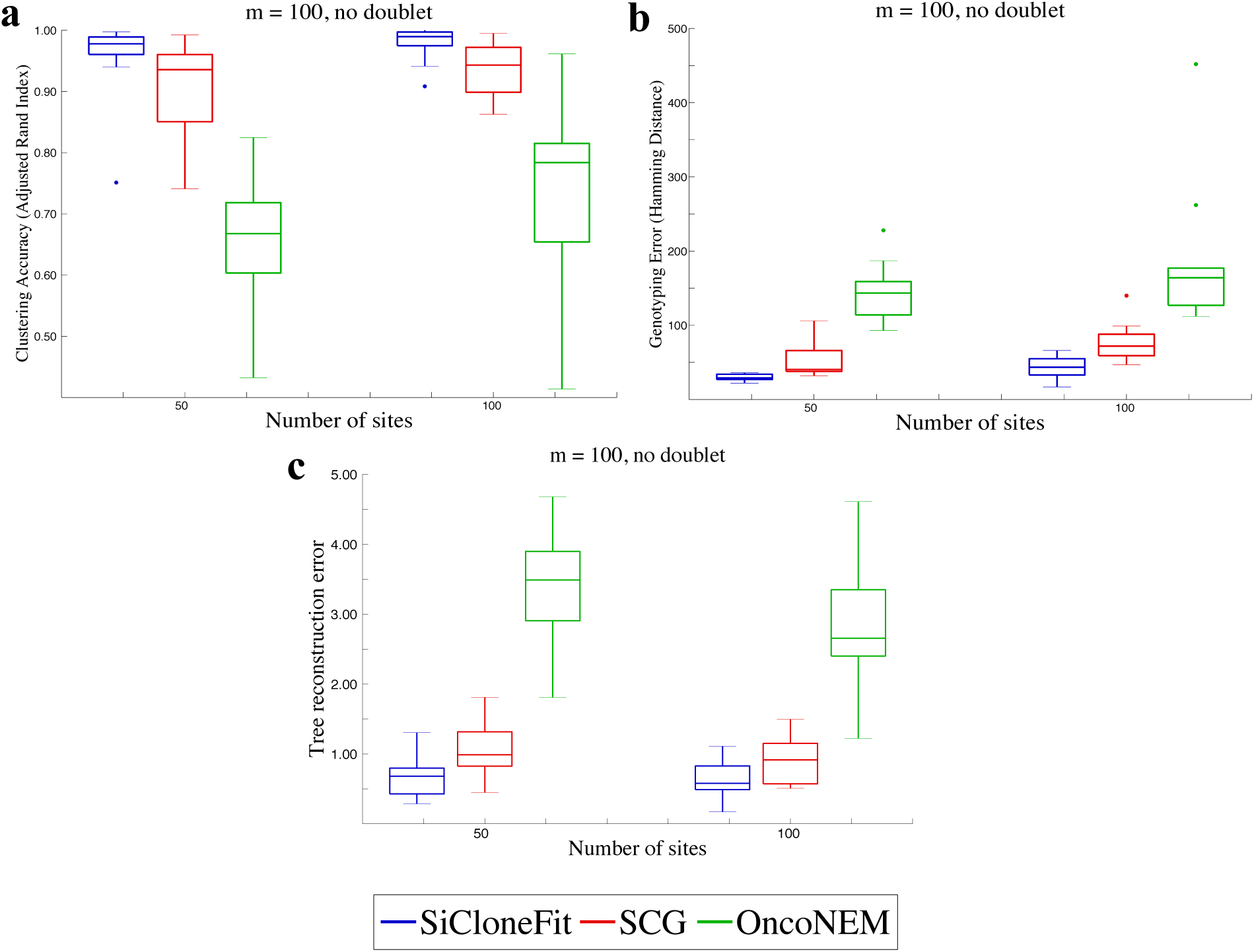
Performance comparison on simulated datasets containing 100 cells. SiCloneFit’s performance is compared against that of SCG and OncoNEM on simulated datasets containing 100 cells for varying numbers of sites. On the x-axis, we have results corresponding to *n* = 50 and *n* = 100. The cells were sampled from *K* = 10 clonal populations. Each box plot summarizes results for 10 simulated datasets with varying clonal phylogeny and varying size of clonal clusters. Comparison of clustering accuracy measured in terms of adjusted rand index that compares the inferred clustering from the ground truth. (b) Comparison based on the genotyping error measured in terms of hamming distance between the true genotype matrix and inferred genotype matrix. (c) Comparison based on the tree reconstruction error measured in terms of pairwise cell shortest-path distance between the true clonal phylogeny and inferred clonal phylogeny.

Next, we performed simulations including 10% doublets to evaluate SiCloneFit’s doublet model. For a fixed number of clones, we simulated datasets with varying number of cells and varying number of sites. SiCloneFit achieved higher clustering accuracy (Supplemental Fig. S12) and genotyping accuracy (Supplemental Fig. S13) compared to SCG. It also achieved lower tree reconstruction error (Supplemental Fig. S14) in all settings except for *m* = 500 and *n* = 100. For some datasets, SCG failed to converge and resulted in low clustering accuracy and high genotyping error. In the presence of higher number of clonal populations, SiCloneFit significantly outperformed SCG (Supplemental Fig. S15). Finally, we tested how inference is affected when missing data and doublets are simultaneously present. Even in the presence of high amount of missing data (15%, 30%), both methods performed well in clustering (Supplemental Fig. S16) the cells to clones, with SiCloneFit performing better than SCG in all settings. For all settings, SiCloneFit’s genotyping (Supplemental Fig. S17) was better than SCG’s. For some datasets, SCG failed to converge and resulted in low clustering accuracy and high genotyping error. SiCloneFit’s tree reconstruction error (Supplemental Fig. S18) was also lower in all but one setting (*n* = 100 and 15% missing data).

### Inference of Clonal Clusters, Genotypes and Phylogeny from Experimental SCS Data

We applied SiCloneFit to two experimental single-cell DNA sequencing datasets from two metastatic colon cancer patients, obtained from the study of Leung *et al.* (Leung et al., 2017). These datasets were generated using a highly-multiplexed single-cell DNA sequencing method (Leung et al., 2016) and a 1000 cancer gene panel was used as the target region for sequencing. These are two of the most recent SCS datasets and contain large numbers of cells and small numbers of mutation sites making the inference difficult.

The first dataset consisted of 178 cells (Leung et al., 2017) obtained from both primary colon tumor and liver metastasis. The original study reported 16 somatic SNVs after variant calling. The reported genotypes were binary values, representing the presence or absence of a mutation at the SNV sites. In the original study, SCITE (Jahn, Kuipers, and Beerenwinkel, 2016) was used for performing phylogenetic analysis of this tumor. However, SCITE operates under the infinite sites assumption and only infers the mutation tree. We ran the four-gamete test on this dataset, which identified 104 (out of 120) pairs of SNV sites violating the four-gamete test indicating potential violation of the infinite sites assumption. After running SiCloneFit on this dataset, we collected the samples from the posterior and computed a maximum clade credibility tree based on the posterior samples as shown in Fig. 3a. Five different clusters were identified from the SiCloneFit posterior samples. The largest cluster (N) consisted of normal cells without any somatic mutation. The primary tumor cells were clustered into two subclones (P1 and P2). Metastatic aneuploid tumor cells were clustered into one subclone (M). There was another cluster (D) consisting of diploid cells (mostly metastatic). The clonal genotype of each cluster was inferred based on the posterior samples. The inferred genotypes are shown in Supplemental Fig. S19. Based on the clonal genotypes, we inferred the ancestral sequences at the internal nodes and this enabled us to find the maximum likelihood solution for placing the mutations on the branches of the clonal phylogeny. First, a heterozygous nonsense mutation was acquired in *APC* along with mutations in *KRAS* oncogene and *TP53* tumor suppressor gene and these initiated the tumor mass. The subclone (D) consisting of diploid cells acquired another mutation in *GATA1* and branched out from the primary tumor mass. The primary tumor subclones developed by acquiring 6 more somatic mutations including a mutation in *CCNE1* oncogene. These mutations were subsequently inherited in the metastatic tumor subclone (M). The accumulation of mutations in *EYS*, *GATA1*, *RBFOX1*, *TRRAP* and *ZNF521* marked the point of metastatic divergence. The two primary tumor subclones were distinguished by the presence/absence of *TPM4* mutation. It was specific to the second primary subclone (P2) and was not identified in any of the tumor cells in the metastasis, suggesting that the first primary subclone (P1) disseminated and established the metastatic tumor mass. We ran MACHINA (El-Kebir, Satas, and Raphael, 2018) on the clonal phylogeny inferred by SiCloneFit for reconstructing the migration history of the tumor clones for this patient. The inferred migration graph (Fig. 3b) had two migrations with comigration number = 1. Since, two anatomical sites were sequenced, the inference of minimum possible comigration number indicates a single-source seeding pattern with colon being the source. The presence of a multi-edge in the migration graph also indicates polyclonal seeding, where liver was seeded by two different clones that originated in colon. However, the first seeding did not result in the clonal expansion, metastatic tumor mass formed after the second seeding that was associated with the mutations in *EYS*, *GATA1*, *RBFOX1*, *TRRAP* and *ZNF521*.

**Figure 3:**
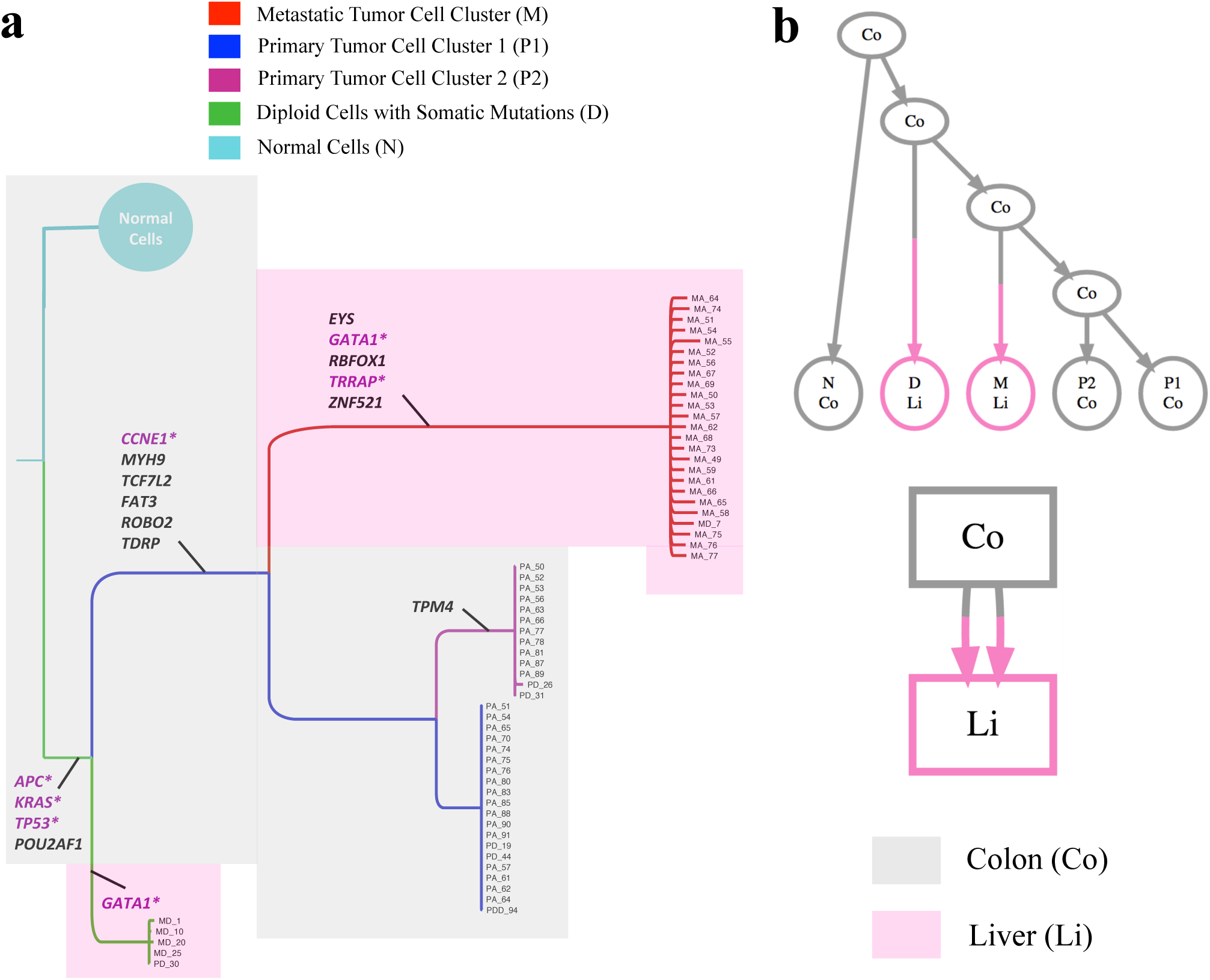
Inference of tumor clones and clonal phylogeny using SiCloneFit on metastatic colorectal cancer patient CRC1. (a) Maximum clade credibility tree reconstructed from the posterior samples obtained using SiCloneFit. Each tumor clone is a cluster of single cells and their genotypes are also inferred. The temporal order of the mutations is reconstructed and mutations are annotated on the branches of the clonal tree. The cancer genes and tumor-suppressor genes are marked in purple. The colors of the shades represent the organ/anatomical site of the origin of the cells. (b) Parsimonious migration history of the tumor clones inferred using MACHINA (El-Kebir, Satas, and Raphael, 2018) with the SiCloneFit inferred clonal tree as input. The top figure shows the clonal tree where the leaves are annotated by the anatomical sites and the anatomical sites annotation of the internal nodes and root are inferred by MACHINA. The bottom figure shows the migration graph of the cells with migration number 2 and comigration number 1. This indicates polyclonal single-source seeding from colon to liver.

For comparison, we ran SCG on this dataset. SCG reported 4 clonal clusters and the inferred clonal genotypes are shown in Supplemental Fig. S20. SCG could not distinguish the primary tumor cells on the basis of the presence/absence of the *TPM4* mutation and genotyped all of them to contain TPM4. Thus it did not report two primary tumor subclones that were detected by SiCloneFit and instead only one primary tumor subclone (all primary tumor cells were assigned to this cluster) was inferred.

The second dataset consisted of 182 cells (Leung et al., 2017) obtained from both primary colon tumor and liver metastasis. The original study reported 36 somatic SNVs after variant calling. The reported genotypes were binary values, representing the presence or absence of a mutation at the SNV sites. After running the four-gamete test on this dataset, we identified 347 (out of 630) pairs of SNV sites violating the four-gamete test indicating potential violation of the infinite sites assumption. After running SiCloneFit on this dataset, we collected the samples from the posterior and computed a maximum clade credibility tree based on the posterior samples as shown in Fig. 4b. Six different clusters were identified in the MPEAR solution based on the posterior samples. The largest cluster (N) consisted of normal cells that did not harbor any somatic mutation. There were two clusters consisting of primary aneuploid tumor cells (P1 and P2) and two clusters consisting of metastatic aneuploid tumor cells (M1 and M2). There was one more cluster (I) comprised of diploid cells that harbored somatic mutations that were completely different from the primary or metastatic clusters, representing an independant clonal lineage consistent with the findings reported by Leung *et al.* (Leung et al., 2017). The clonal genotype of each cluster was inferred based on the posterior samples. The inferred genotypes are shown in Supplemental Fig. S21. Based on the clonal genotypes, we inferred the ancestral sequences at the internal nodes and this enabled us to find the maximum likelihood solution for placing the mutations on the branches of the clonal phylogeny. The first primary tumor clone (P1) evolved from the normal cells by acquiring 8 mutations including mutations in *APC*, *NRAS*, *CDK4* and *TP53*. After that 4 additional mutations (*CHN1, APC, LINGO2, IL21R*) were acquired before the first metastatic cluster (M1) diverged. After dissemination into liver, the first metatstatic subclone (M1) continued to evolve and acquired a number of metastasis-specific mutations (e.g., *SPEN*, *IL7R*, *PIK3CG*, *F8*, *LINGO2*). Before the divergence of the second metastatic subclone (M2), two more mutations (*FHIT, ATP7B*) were acquired that were also present in the second primary tumor subclone P2. The second primary tumor clone (P2) acquired two additional mutations in *LRP1B* and *LINGO2* that were not present in either metastatic clone. The second metastatic clone disseminated after acquiring the *ATP7B* mutation and further expanded the liver tumor mass by acquiring 7 additional mutations (e.g., *PTPRD*, *NR4A3*, *HELZ*, *TSHZ3*). In the original study, SCITE identified two different lineages for metastatic cells and inferred 4 mutations (*FHIT*, *ATP7B*, *APC*, and *CHN1*) between the two metastatic divergence events. These were called as “bridge mutations”. However, statistical analysis (Leung et al., 2017) of the four bridge mutations provided strong evidence only for two of them (*FHIT* and *ATP7B*), and the placement of the other two bridge mutations were uncertain. In our analysis, SiCloneFit correctly identified the two strongly supported bridge mutations (*FHIT* and *ATP7B*) as the mutations between the two metastatic divergence events. The other two putative bridge mutations were identified as non-bridge and placed before the divergence of first metastatic subclone. Other than the precursor mutations shared with the primary tumor clones, the metastatic tumor clones had three more mutations in common (*PTPRD*, *FUS* and *LINGO2*). This is an evidence for a potential convergent evolution. To evaluate the accuracy of this, we performed the mixture-model Bayesian binomial test (Leung et al., 2017), which provided strong evidence of recurrence for two of these mutations (*FUS* and *LINGO2*, see Supplemental Fig. S22, Supplemental Results for details). Apart from the primary and metastatic tumor clones, there was another cluster (I) consisting of 7 primary diploid cells that had completely independent somatic mutations. These cells acquired mutations in *SPEN, ALK, ATR, NR3C2* and *EPHB6* but did not share any other mutations with the primary or metastatic tumor cells, representing an entirely different tumor lineage that did not expand significantly. We reconstructed the migration history of the tumor clones by running MACHINA (El-Kebir, Satas, and Raphael, 2018) on the SiCloneFit inferred clonal phylogeny, whose leaves (clonal clusters) were annotated by the anatomical site of origin of the associated cells. The inferred migration graph (Fig. 4b) had two migrations with comigration number = 1 (also the minimum possible comigration number for two anatomical sites), indicating polyclonal single-source seeding from colon to liver. Here, both the seeding events led to expansion of tumor mass in liver and resulted in two different metastatic subclones.

**Figure 4:**
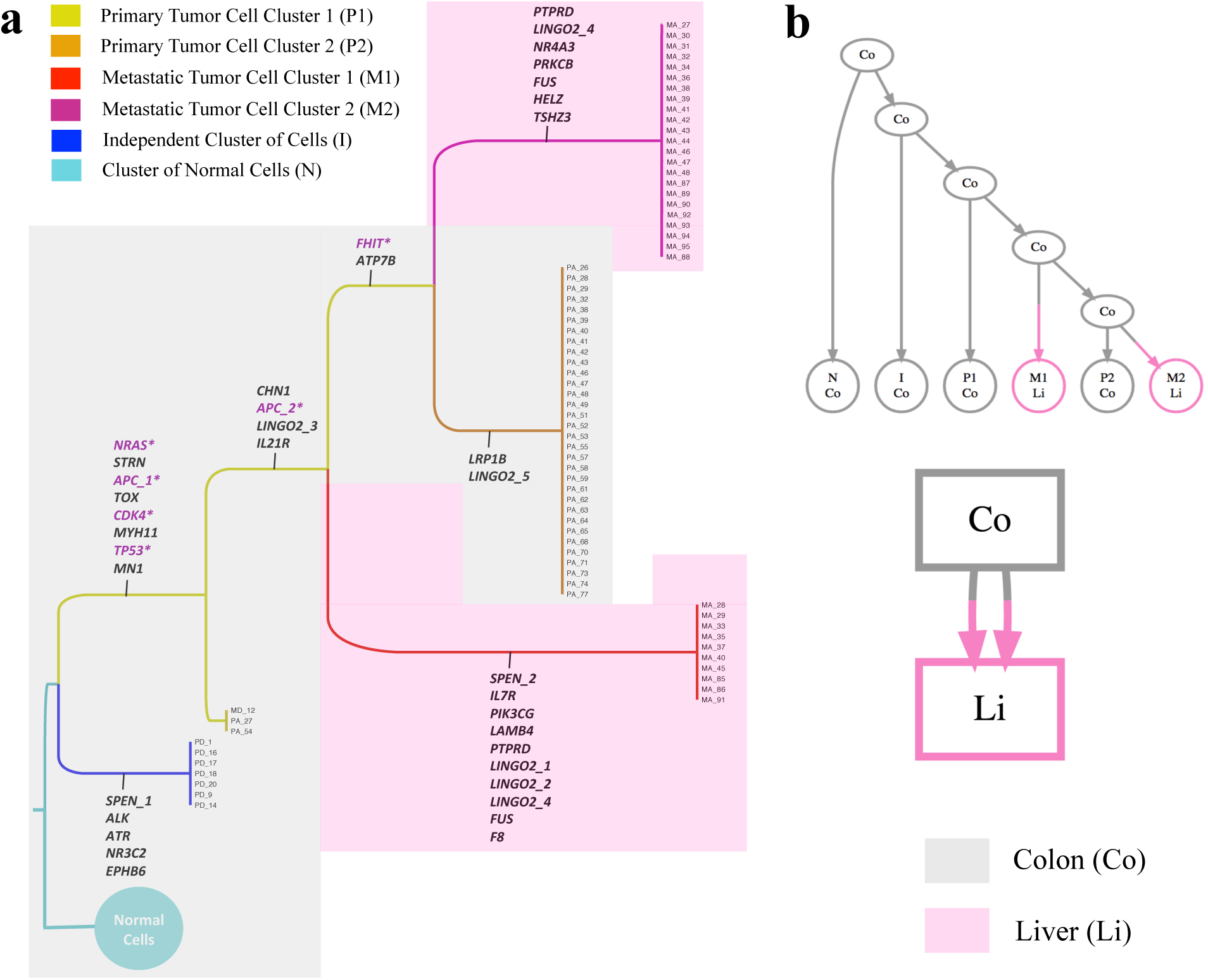
Inference of tumor clones and clonal phylogeny using SiCloneFit for metastatic colorectal cancer patient CRC2. (a) Maximum clade credibility tree reconstructed from the posterior samples obtained using SiCloneFit. Each tumor clone is a cluster of single cells and their genotypes are also inferred. The temporal order of the mutations is reconstructed and mutations are annotated on the branches of the clonal tree. The cancer genes and tumor-suppressor genes are marked in purple. The colors of the shades represent the organ/anatomical site of the origin of the cells. (b) Parsimonious migration history of the tumor clones inferred using MACHINA (El-Kebir, Satas, and Raphael, 2018) with the SiCloneFit inferred clonal tree as input. The top figure shows the clonal tree where the leaves are annotated by the anatomical sites and the anatomical sites annotation of the internal nodes and root are inferred by MACHINA. The bottom figure shows the migration graph of the cells with migration number 2 and comigration number 1. This indicates polyclonal single-source seeding from colon to liver.

SCG reported 5 clonal clusters from this dataset (Supplemental Fig. S23). Clustering and genotyping results of SCG mostly agreed with that of SiCloneFit. However, SCG failed to detect two primary tumor subclones and instead clustered them together into one subclone resulting in incorrect genotyping for those cells. Furthermore, SCG did not infer any phylogeny.

## Discussion

Inference of tumor subclones and their evolutionary history is of paramount importance given their contribution to drug resistance and therapeutic relapse. While this problem has been investigated in depth in the context of bulk sequencing data, methods are lacking for SCS data, most promising and high-resolution data for studying tumor heterogeneity. Here, we reported on SiCloneFit, a novel probabilistic framework for inferring the number and structure of tumor clones, their genotypes and evolutionary history from noisy somatic SNV profiles of single cells. Our unified framework jointly reconstructs the tumor clones as clusters of single cells as well as their genealogical relationship in the form of a clonal phylogeny. In this process, SiCloneFit accounts for the effects of mutational events (point mutations, LOH, deletion) in the evolutionary history of the tumor via a finite-sites model of evolution and denoises the effects of technical artifacts such as allelic dropout, false-positive errors, missing entries and cell-doublets to infer the clonal genotypes. SiCloneFit employs a Gibbs sampling algorithm consisting of partial reversible-jump MCMC, partial Gibbs updates for estimating the latent variables by sampling from the posterior distribution. A major distinguishing feature of SiCloneFit is that it jointly solves the subclonal reconstruction and tumor phylogeny inference problems from SCS datasets whereas existing methods either cluster the cells into subclones or infers a tumor phylogeny. The phylogeny inference methods (except SiFit) also rely on infinite sites assumption to restrict the search space. On the contrary, SiCloneFit employs a finite-sites model of evolution to account for mutation recurrence and losses. At the same time, SiCloneFit accounts for cell doublets, an important technical artifact that is not dealt with by existing single-cell phylogeny inference methods.

We assessed SiCloneFit’s performance through a comprehensive set of simulation studies aimed at creating experimental settings corresponding to different aspects of modern SCS datasets. Datasets were generated with a varying number of cells, genomic sites and tumor subclones, a wide range of error rates, varying amount of missing data and cell doublets. In simulated benchmarks, SiCloneFit outperformed the state-of-the-art methods based on different metrics for evaluating its performance in inferring the clonal clusters, clonal genotypes and the clonal evolutionary history. We also applied SiCloneFit on two targeted SCS datasets from two metastatic colon cancer patients for studying the intratumor heterogeneity. For these tumors, SiCloneFit inferred the primary and metastatic subclones as clusters of single cells, inferred their genotypes, reconstructed the genealogy of these subclones and inferred the temporal order of the mutations in their evolutionary history revealing mutations that potentially played an important role in metastatic divergence.

SiCloneFit’s model is flexible, and more complex model of evolution can be incorporated to account for copy number information. The inclusion of copy number variations along with SNVs can improve the estimation of subclones and help in uncovering the interaction between SNV and CNVs at the single cell level. The error model can be further extended to utilize reference and variant read counts at each mutation site in each cell as the input data instead of presence/absence of mutation inferred by a variant caller.

In closing, SiCloneFit advances the understanding of intratumor heterogeneity and clonal evolution through improved computational analysis of SCS data. As SCS becomes more high-throughput generating somatic SNV profiles for thousands of cells, SiCloneFit will be very helpful in reconstructing the tumor clones and clonal phylogeny from such large datasets. Being capable of handling doublets, SiCloneFit will find important applications in removing doublets, as their percentage can be high in more high-throughput datasets. Methods like SiCloneFit will have important translational applications for improving cancer diagnosis, treatment and therapy in clinical applications.

## Methods

### Model Description

We assume that we have measurements from *m* single cells. For each cell, *n* somatic single nucleotide variant (SNV) sites have been measured. The data can be represented by a matrix *D*_*n×m*_ = (*D*_*ij*_) of observed genotypes, where *D*_*ij*_ is the observed genotype at the *i*^*th*^ site of cell *j*. Let *g*_*t*_ be the set of possible true genotype values for the SNVs, and *g*_*o*_ be the set of observable values for the SNVs. For binary measurements for SNVs, *g*_*t*_ = {0, 1}, whereas *g*_*o*_ = {0, 1*, X*}, where 0, 1 and *X* denote the absence of mutation, presence of mutation, and missing value respectively. If ternary measurements are available for SNVs, *g*_*t*_ = {0, 1, 2} and *g*_*o*_ = {0, 1, 2*, X*}, where 0 denotes homozygous reference genotype, 1 and 2 denote heterozygous, and homozygous non-reference genotypes, respectively, and *X* denotes missing data.

We assume that there is a set of *K* clonal populations from which *m* single cells are sampled and the clonal populations can be placed at the leaves of a clonal phylogeny, *𝒯*. Each clonal population consists of a set of cells that have identical genotype (with respect to the set of mutations in consideration) and a common ancestor. The genotype vector associated with a clone *c* is called clonal genotype (denoted by *G*_*c*_) and it records the genotype values for all *n* sites for the corresponding clone. The true genotype vector of each cell is identical to the clonal genotype of the clonal population where it belongs to. The clonal genotype matrix, *G*_*K×n*_, represents the clonal genotypes of *K* clones. It is important to note that, *K*, the number of clones is unknown. To automatically infer the number of clones and assign the cells to clones, we introduce a tree-structured infinite mixture model. (Meeds et al., 2008) describes a nonparametric Bayesian prior over trees similar to mixture models using a Chinese restaurant process (CRP) (Pitman, 2006) prior. For this tree-structured CRP, each node of the tree represents a cluster. In our model, we extend this idea to define a nonparametric Bayesian prior over binary trees, leaves of which represent the mixture components (clonal clusters). A Chinese restaurant process defines a distribution for partitioning customers into different tables. In our problem, single cells are analogous to customers and clonal clusters are analogous to tables. Let *c*_*j*_ denote the cluster assignment for cell *j* and assume that cells 1: *j -* 1 have already been assigned to clonal clusters {1, …, |***c***_1:*j-*1_*|*}, where *|****c***_1:*j-*1_*|* denotes the number of clusters induced by the cluster indicators of *j -* 1 cells. The cluster assignment of cell *j*, *c*_*j*_ is based on the distribution defined by a Chinese restaurant process is given by

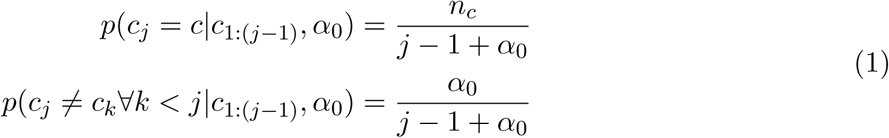

where *n*_*c*_ denotes the number of cells already assigned (excluding cell *j*) to cluster *c*. *α*_0_ is the concentration parameter for the CRP model.

The clonal phylogeny, *𝒯*, is a rooted directed binary tree whose number of leaves is equal to the number of clonal clusters, *K* = *|****c****|* defined by the assignment of *m* cells to different clusters by the CRP. The root of *𝒯* represents normal (unmutated) genotype and somatic mutations are accumulated along the branches of the phylogeny. Each leaf in the clonal phylogeny corresponds to a clonal cluster, *c ∈* {1, …, *K*} and is associated with a clonal genotype *G*_*c*_ that records the set of mutations accumulated along the branches from the root. To model the evolution of the clonal genotypes, we employ a finite-site model of evolution, *ℳ*_*λ*_, that accounts for the effects of point mutations, deletion and loss of heterozygosity on the clonal genotypes. The model of evolution assigns transition probabilities to different genotype transitions along the branches of the clonal phylogeny. The true genotype of each cell is identical to the clonal genotype of the clonal cluster where it is assigned. However, observed genotypes of single cells differ from their true genotype due to amplification errors introduced during the single-cell sequencing work flow. The effect of amplification errors is modeled using an error model distribution parameterized by FP error rate, *α* and FN error rate, *β*. The generative process can be described as follows:

1. draw *α*_0_ *∼ Gamma*(*a, b*), *α ∼ Beta*(*a*_*α*_*, b*_*α*_), *β ∼ Beta*(*a*_*β*_*, b*_*β*_)
2. For *j ∈* {1, 2, …, *m*}, draw *c*_*j*_ *∼ CRP* (*α*_0_). From this, derive *K* = *|****c****|*, the total number of clusters (or clones) implicitly defined by ***c***.
3. draw *𝒯 ∼ T*_*prior*_(*K*).
4. For *λ ∈ ℳ*_*λ*_, draw 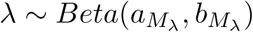
5. For *κ ∈* {1, 2, …, *K*}, draw *G*_*κ*_ *∼ F* (*G*_*κ*_*| 𝒯, ℳ* _*λ*_).
6. For *j ∈* {1, 2, …, *m*} and *i ∈ {*1, 2, …, *n*}, draw 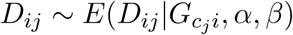.

***c*** denotes the clonal assignments of all cells. *T*_*prior*_ is the prior distribution on phylogenetic trees for a fixed number of leaves. *ℳ*_*λ*_ denotes the set of parameters in the finite-sites model of evolution. *F* denotes a distribution on the genotypes at the leaves of a phylogenetic tree and can be computed using Felsenstein’s pruning algorithm Felsenstein, 1981 given the phylogeny and a finite-site model of evolution. *E* is the error model distribution that relates the observed genotype at locus *i* for cell *j*, *D*_*ij*_ to clonal genotype 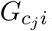. *a, b, a*_*α*_*, b*_*α*_*, a*_*β*_*, b*_*β*_*, a*_*M*_ *, b*_*M*_ denote different hyperparameters used in this model.

In addition to the variables in the singlet model, the doublet-aware model of SiCloneFit incorporates a Bernoulli variable for each cell for indicating whether the cell is a singlet or a doublet, a Beta distributed variable for doublet rate and a secondary cluster indicator for each cell so that a doublet can be assigned to two clonal clusters. Supplemental Table S6 defines the expected genotype for a doublet. These additional variables are described in Supplemental Table S7. The generative process for the doublet-aware model is described in detail in Supplemental Methods.

### Model of Evolution and Error Model

To capture the effect of point mutations, LOH and deletion on the clonal genotypes along the branches of clonal phylogeny, we employ a finite-site model of evolution similar to the one introduced in SiFit (Zafar et al., 2017). The finite-site model of evolution, *ℳ*_*λ*_, is modeled using a continuous-time Markov chain that assigns a probability with each possible transition of genotypes. The transition rate matrix of the continuous-time Markov chain for binary and ternary genotypes can be defined based on branch length, *t*, and parameters *λ*_*r*_ and *λ*_*l*_, accounting for the effects of recurrent mutation and mutation loss, respectively. These are described in detail in Supplemental Methods.

To account for FP and FN errors in SCS data, we introduce an error model distribution, 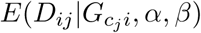, which gives the probability of observing genotype *D*_*ij*_ for locus *i* in cell *j*, given the true clonal genotype 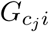. The error model distribution for ternary and binary data are shown in Supplemental Table 4 and Supplemental Table 5 respectively.

### Posterior Distribution

The posterior distribution *𝒫* over the latent variables of the SiCloneFit model is given by

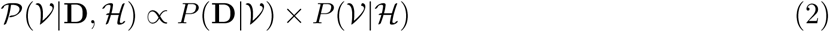

where *𝒱* denotes the set of latent variables in the model, *𝒱* = {**c**, **G***, 𝒯, ℳ*_*λ*_*, α, β, α*_0_}. **c** is a vector containing cluster assignment for all cells and it implicitly defines the number of clones *K*. *ℋ* is the set of fixed hyper-parameters of the model, *ℋ* = {*a*_*α*_*, b*_*α*_*, a*_*β*_*, b*_*β*_*, a*_*M*_ *, b*_*M*_ *, a, b*}. In Eq. (2), the term *P* (**D***| 𝒱*) denotes the likelihood of the model and the term *P* (*𝒱| ℋ*) denotes the product of prior probabilities. The posterior distribution for the doublet-aware model is described in Supplemental Methods.

### Likelihood Function

The likelihood function employed by SiCloneFit is given by

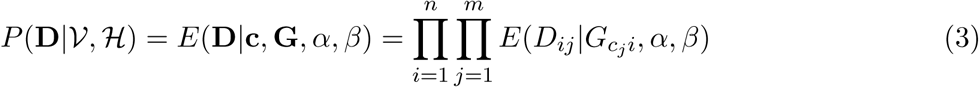

In Eq. (3), 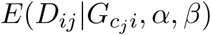 is given by the error model distribution of observing genotype *D*_*ij*_ for site *i* in cell *j*, given the true clonal genotype 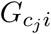 and is parameterized by *α* and *β*. This error model is based on the error model of SiFit (Zafar et al., 2017). The likelihood of the doublet-aware model is described in Supplemental Methods.

### Prior Distributions

The SiCloneFit model incorporates a compound prior given by

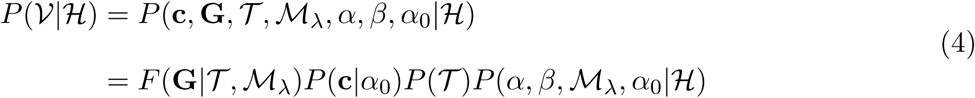

Where

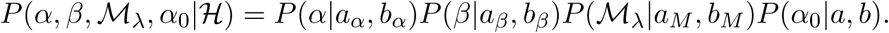

*F* (**G***| 𝒯, ℳ*_*λ*_) denotes the prior distribution on the clonal genotype matrix given a clonal phylogeny *𝒯* and parameters of the model of evolution *ℳ*_*λ*_, and it can be efficiently calculated using Felsenstein’s pruning algorithm (Felsenstein, 1981) assuming sites are independent and identically distributed. *P* (**c***|α*_0_) denotes the prior probability of partitioning *m* single cells into *K* (*K* is the number of clusters defined by **c**) clusters under a CRP with concentration parameter *α*_0_. *P* (*𝒯*) denotes the prior probability on the clonal phylogeny. This is a product of prior on topology and prior on branch length. We consider uniform distribution for the prior on topology and exponential distribution for the prior on branch lengths. As the values of the error rate parameters *α*, *β* and the parameters of the model of evolution *ℳ*_*λ*_ lie between 0 and 1, we use Beta distribution as their prior. For the concentration parameter *α*_0_, we assume a Gamma prior as suggested in (Escobar and West, 1995). We set the value of hyperparameters for the Gamma distribution to *a* = 1 and *b* = 1 for all the analyses performed, but these are user-specified parameters in the software. All the prior distributions are described in detail in Supplemental Methods. The doublet-aware model of SiCloneFit contains additional parameters for indicating whether a cell is a singlet or a doublet, doublet rate and assigning a cell to two clonal clusters and the associated prior distributions are described in Supplemental Methods.

### Inference

We designed a Markov chain Monte Carlo (MCMC) sampling procedure based on the Gibbs sampling algorithm to estimate the latent variables according to Eq. (2). Our algorithm is inspired by a partial Metropolis-Hastings, partial Gibbs sampling algorithm described in (Neal, 2000). In each iteration, the sampler first samples new cluster indicators, **c**^*\**^, for all the cells using partial Metropolis-Hastings partial Gibbs updates. During this, the dimensionality of the sample may change due to addition of a new cluster (resulting in addition of new edges in the clonal phylogeny) or removal of an existing singleton cluster (resulting in removal of existing edges from the clonal phylogeny). In case the dimensionality changes, the absolute value of the determinant of the Jacobian matrix is also taken into account, which results in partial reversible-jump MCMC (Green, 1995) updates. When such dimension changing moves are accepted, the corresponding new clonal phylogeny *𝒯 \** and new clonal genotype matrix **G**^*\**^ are also accepted. The sampler next samples a new clonal phylogeny and new parameters of the model of evolution with the help of a Metropolis-Hastings MCMC sampler. After that, new clonal genotype for each clonal cluster is sampled from the conditional posterior distribution. To sample new values of the error rate parameters from their corresponding conditional posterior distributions, our sampler employs rejection sampling. Finally, the concentration parameter *α*_0_ is sampled based on the method described in (Escobar and West, 1995). The sampling algorithms for both the singlet and doublet models of SiCloneFit are described in detail in Supplemental Methods.

### Software Availability

SiCloneFit has been implemented in Java and is freely available at https://bitbucket.org/hamimzafar/siclonefit, released under the MIT license.

## Acknowledgments

L.N. was supported by National Science Foundation Grant IIS-1812822. K.C. was supported by the National Cancer Institute (grant R01 CA172652), a Cancer Center support grant (P30 CA016672), and the Andrew Sabin Family Foundation.

## References

Amigó, Enrique et al. (2009). “A Comparison of Extrinsic Clustering Evaluation Metrics Based on Formal Constraints”. In: Inf. Retr. 12.4, pp. 461–486.

Baslan, Timour and James Hicks (2017). “Unravelling biology and shifting paradigms in cancer with single-cell sequencing”. In: Nat Rev Cancer 17.9. Perspectives, pp. 557–569.

Burrell, Rebecca A. et al. (2013). “The causes and consequences of genetic heterogeneity in cancer evolution”. In: Nature 501.7467, pp. 338–345.

Davis, Alexander and Nicholas E. Navin (2016). “Computing tumor trees from single cells”. In: Genome Biology 17.1, pp. 1–4.

Deshwar, Amit G. et al. (2015). “PhyloWGS: Reconstructing subclonal composition and evolution from whole-genome sequencing of tumors”. In: Genome Biology 16.1, pp. 1–20.

El-Kebir, Mohammed, Gryte Satas, and Benjamin J. Raphael (2018). “Inferring parsimonious migration histories for metastatic cancers”. In: Nature Genetics 50.5, pp. 718–726.

El-Kebir, Mohammed et al. (2016). “Inferring the Mutational History of a Tumor Using Multi-state Perfect Phylogeny Mixtures”. In: Cell Systems 3.1, pp. 43–53.

Escobar, Michael D. and Mike West (1995). “Bayesian Density Estimation and Inference Using Mixtures”. In: Journal of the American Statistical Association 90.430, pp. 577–588.

Felsenstein, Joseph (1981). “Evolutionary trees from DNA sequences: A maximum likelihood approach”. In: Journal of Molecular Evolution 17.6, pp. 368–376.

Fritsch, Arno and Katja Ickstadt (2009). “Improved criteria for clustering based on the posterior similarity matrix”. en. In: Bayesian Anal. 4.2, pp. 367–391.

Gawad, Charles, Winston Koh, and Stephen R. Quake (2014). “Dissecting the clonal origins of childhood acute lymphoblastic leukemia by single-cell genomics”. In: Proceedings of the National Academy of Sciences 111.50, pp. 17947–17952.

Gerlinger, Marco et al. (2014). “Genomic architecture and evolution of clear cell renal cell carcinomas defined by multiregion sequencing”. In: Nature Genetics 46.3. Article, pp. 225–236.

Gerlinger, Marco et al. (2012). “Intratumor Heterogeneity and Branched Evolution Revealed by Multiregion Sequencing”. In: New England Journal of Medicine 366.10, pp. 883–892.

Gillies, Robert J., Daniel Verduzco, and Robert A. Gatenby (2012). “Evolutionary dynamics of carcinogenesis and why targeted therapy does not work”. In: Nat Rev Cancer 12.7, pp. 487–493.

Green, Peter J. (1995). “Reversible jump Markov chain Monte Carlo computation and Bayesian model determination”. In: Biometrika 82.4, pp. 711–732.

Hou, Yong et al. (2012). “Single-Cell Exome Sequencing and Monoclonal Evolution of a JAK2-Negative Myeloproliferative Neoplasm”. In: Cell 148.5, pp. 873 –885.

Jahn, Katharina, Jack Kuipers, and Niko Beerenwinkel (2016). “Tree inference for single-cell data”. In: Genome Biology 17.1, pp. 1–17.

Jiang, Yuchao et al. (2016). “Assessing intratumor heterogeneity and tracking longitudinal and spatial clonal evolutionary history by next-generation sequencing”. In: Proceedings of the National Academy of Sciences 113.37, E5528–E5537.

Kandoth, Cyriac et al. (2013). “Mutational landscape and significance across 12 major cancer types”. In: Nature 502.7471. Article, pp. 333–339.

Kuipers, Jack et al. (2017). “Single-cell sequencing data reveal widespread recurrence and loss of mutational hits in the life histories of tumors”. In: Genome Research 27.11, pp. 1885–1894.

Landau, Dan A. et al. (2015). “Mutations driving CLL and their evolution in progression and relapse”. In: Nature 526.7574. Article, pp. 525–530.

Leung, Marco L. et al. (2016). “Highly multiplexed targeted DNA sequencing from single nuclei”. In: Nat. Protocols 11.2. Protocol, pp. 214–235.

Leung, Marco L. et al. (2017). “Single-cell DNA sequencing reveals a late-dissemination model in metastatic colorectal cancer”. In: Genome Research.

Li, Yingrui et al. (2012). “Single-cell sequencing analysis characterizes common and cell-lineage-specific mutations in a muscle-invasive bladder cancer”. In: GigaScience 1.1, p. 12.

Meeds, E. W. et al. (2008). “Learning stick-figure models using nonparametric Bayesian priors over trees”. In: 2008 IEEE Conference on Computer Vision and Pattern Recognition, pp. 1–8.

Merlo, Lauren M.F. et al. (2006). “Cancer as an evolutionary and ecological process”. In: Nat Rev Cancer 6.12, pp. 924–935.

Navin, Nicholas (2014). “Cancer genomics: one cell at a time”. In: Genome Biology 15.8, pp. 452–465.

Navin, Nicholas E. (2015). “The first five years of single-cell cancer genomics and beyond”. In: Genome Res 25.10, pp. 1499–1507.

Neal, Radford M. (2000). “Markov Chain Sampling Methods for Dirichlet Process Mixture Models”. In: Journal of Computational and Graphical Statistics 9.2, pp. 249–265.

Nik-Zainal, Serena et al. (2012). “The Life History of 21 Breast Cancers”. In: Cell 149.5, pp. 994–1007.

Nowell, PC (1976). “The clonal evolution of tumor cell populations”. In: Science 194.4260, pp. 23–28.

Pepper, John W. et al. (2009). “SYNTHESIS: Cancer research meets evolutionary biology”. In: Evolutionary Applications 2.1, pp. 62–70.

Pitman, Jim (2006). “Combinatorial Stochastic Processes”. In: Ecole dEt de Probabilits de Saint-Flour XXXII.

Ross, Edith M. and Florian Markowetz (2016). “OncoNEM: inferring tumor evolution from single-cell sequencing data”. In: Genome Biology 17.1, pp. 1–14.

Roth, Andrew et al. (2016). “Clonal genotype and population structure inference from single-cell tumor sequencing”. In: Nat Meth 13.7. Brief Communication, pp. 573–576.

Roth, Andrew et al. (2014). “PyClone: statistical inference of clonal population structure in cancer”. In: Nat Meth 11.4, pp. 396–398.

Sainudiin, Raazesh and Amandine Veber (2016). “A Beta-splitting model for evolutionary trees”. In: Royal Society Open Science 3.

Schliep, K.P. (2011). “phangorn: phylogenetic analysis in R”. In: Bioinformatics 27.4, pp. 592–593.

Shah, Sohrab P. et al. (2012). “The clonal and mutational evolution spectrum of primary triple-negative breast cancers”. In: Nature 486.7403, pp. 395–399.

Sukumaran, Jeet and Mark T. Holder (2010). “DendroPy: a Python library for phylogenetic computing”. In: Bioinformatics 26.12, pp. 1569–1571.

Turke, Alexa B. et al. (2010). “Preexistence and Clonal Selection of MET Amplification in EGFR Mutant NSCLC”. In: Cancer Cell 17.1, pp. 77–88.

Vogelstein, Bert et al. (2013). “Cancer Genome Landscapes”. In: Science 339.6127, pp. 1546–1558.

Wang, Yong et al. (2014). “Clonal evolution in breast cancer revealed by single nucleus genome sequencing”. In: Nature 512.7513, pp. 155–160.

Wu, Xiaochong et al. (2012). “Clonal selection drives genetic divergence of metastatic medulloblastoma”. In: Nature 482, 529 EP –.

Yates, Lucy R. and Peter J. Campbell (2012). “Evolution of the cancer genome”. In: Nat Rev Genet 13.11, pp. 795–806.

Yates, Lucy R. et al. (2015). “Subclonal diversification of primary breast cancer revealed by multiregion sequencing”. In: Nat Med 21.7. Article, pp. 751–759.

Zafar, Hamim et al. (2018). “Computational approaches for inferring tumor evolution from single-cell genomic data”. In: Current Opinion in Systems Biology 7. Future of systems biology Genomics and epigenomics, pp. 16 –25.

Zafar, Hamim et al. (2017). “SiFit: inferring tumor trees from single-cell sequencing data under finite-sites models”. In: Genome Biology 18.1, p. 178.

